# The role of changes in environmental quality in multitrait plastic responses to environmental and social change in the model microalga *Chlamydomonas reinhardtii*

**DOI:** 10.1101/2020.12.16.422493

**Authors:** Ignacio J. Melero-Jiménez, Antonio Flores-Moya, Sinéad Collins

**Affiliations:** Departamento de Botánica y Fisiología Vegetal, Facultad de Ciencias, Universidad de Málaga, Málaga, Spain; Institute of Evolutionary Biology, School of Biological Sciences, University of Edinburgh, Edinburgh, UK

**Keywords:** *Chlamydomonas reinhardtii*, environmental quality, intraspecific variation, maximum growth rate, carbon dioxide, temperature, light, carbon use efficiency, photosynthesis, reactive oxygen, plasticity

## Abstract

Intraspecific variation plays a key role in species’ responses to environmental change; however, little is known about the role of changes in environmental quality (the population growth rate an environment supports) on intraspecific trait variation. Here, we hypothesize that intraspecific trait variation will be higher in ameliorated environments than in degraded ones. We first measure the range of multitrait phenotypes over a range of environmental qualities for three strains and two evolutionary histories of *Chlamydomonas reinhardtii* in laboratory conditions. We then explore how environmental quality and trait variation affect the predictability of lineage frequencies when lineage pairs are grown in indirect co-culture. Our results show that environmental quality has the potential to affect intraspecific variability both in terms of the variation in expressed trait values, and in terms of the genotype composition of rapidly growing populations. We found low phenotypic variability in degraded or same-quality environments and high phenotypic variability in ameliorated conditions. This variation can affect population composition, as monoculture growth rate is a less reliable predictor of lineage frequencies in ameliorated environments. Our study highlights that understanding whether populations experience environmental change as an increase or a decrease in quality relative to their recent history affects the changes in trait variation during plastic responses, including growth responses to the presence of conspecifics. This points towards a fundamental role for changes in overall environmental quality in driving phenotypic variation within closely-related populations, with implications for microevolution.

## 1. INTRODUCTION

Environmental change can involve both amelioration and deterioration of environments from the point of view of organisms (Howes et al., 2015). Here, we define ameliorated environments are those that allow population growth rate to increase, while degraded ones cause population growth rates to decrease. There is mounting evidence that there is variation among closely-related lineages (such as might form a monospecific population or bloom) in how they respond to environmental change (Batista et al., 2018; Boyd et al., 2013; Schwaderer et al., 2011; Shan et al., 2019). Since phytoplankton are the base of aquatic ecosystems (Kirk, 1994), understanding how trait variation (as cell size, photosynthesis-related traits, cell division rates, and growth response to conspecifics) is affected by different kinds of environmental shifts is crucial to understanding effects on higher trophic levels and nutrient cycling. It is projected that species or lineages that respond to environmental change by increasing population growth rates will become more dominant in ecosystems (Bartosiewicz et al., 2019; Ma et al., 2019; Visser et al., 2016). However, many studies that explore the responses of phytoplankton to environmental change focus on degraded environments where population growth is reduced, sometimes severely (Collins, 2016). This motivated us to test how environmental changes that encompass both amelioration and degradation affect intraspecific trait variation. Our rationale for categorizing environments based on their effect on population growth rate is that this allows a meaningful comparison of qualitatively different environments that can be easily linked to effects on short-term eco-evolutionary dynamics, such as genotype sorting. While one can simply use growth rate to rank environments, we find in our study that it is the change in growth rate relative to the ancestral environment that appears important; so the same environment may represent an ameliorated environment for one population, and a same-quality environment for another.

Figure 1 shows a schematic of three possible trait variation-environmental quality relationships (Figure 1). To include multiple traits that can covary in a number of ways, we define multitrait phenotypes to be phenotypes made up of two or more traits where trait values and trait correlations can differ between populations. We reason that in extremely poor quality environments, the number of multitrait phenotypes expressed would be lower, since very few trait combinations will allow populations to escape extinction by dilution in batch culture experiments, or have a non-negative population growth rate under other conditions. However, in higher quality environments, a higher number of trait combinations would allow populations to have non-negative growth rates.

**Figure 1.**
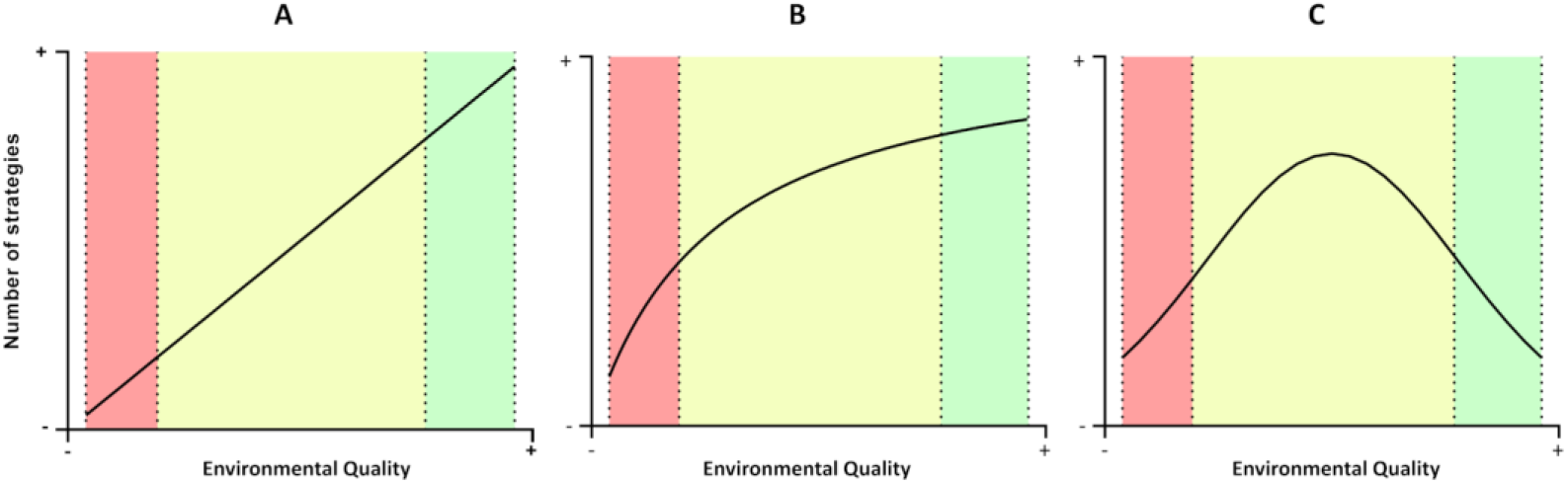
Diagrams showing three possible patterns of the relationship between environmental quality and number of multitrait phenotypes. (A): The number of multitrait phenotypes increases when environmental quality increases. (B): The number of multitrait phenotypes increases when the environmental quality increases until reaching a maximum number of strategies. (C): The number of multitrait phenotypes has a maximum in intermediate-quality environments

Here, we explore the effect of environmental quality on multitrait phenotypes. First, we show that *Chlamydomonas reinhardtii* has a different range of multitrait phenotypes (growth rate, photosynthetic rate, chlorophyll autofluorescence, size cell, and <u>r</u>eactive <u>o</u>xygen <u>s</u>pecies (ROS) production) in environments that differ in terms of quality. In particular, we find that multitrait variability is is higher in ameliorated conditions than elsewhere. Thus, changes in environmental quality have the potential to affect intraspecific trait variability. Second, we explore whether differences in trait variability across abiotic environments affects responses to biotic environments by examining the relationship between maximum population growth rate in monoculture and in indirect co-culture with conspecifics across a range of environmental qualities. We find that during exponential growth, environmental quality can affect the predictability of lineage frequencies during co-culture, but that this depends on the evolutionary history of the populations. Finally, we discuss the potential implications of our results in terms of the potential for rapid evolution by genotype sorting following abrupt environmental shifts.

## 2. MATERIAL AND METHODS

### 2.1 Experimental organism and culture conditions

The strains CC-125, CC-1691, CC-2931 of *Chlamydomonas reinhardtii* were used in these experiments. We used single-strain populations that were previously evolved in ambient CO_2_ and high CO_2_ (2000 μatm) for 90 growth cycles in TAP media under similar measured temperature and light conditions that we use here for our control HL conditions (Lindberg & Collins, 2020). For each strain, we used 3 independently-evolved high CO_2_ populations, and 3 independently-evolved ambient CO_2_ populations, for a total of 18 independently-evolved populations in this study. We refer to populations evolved at high CO_2_ as “high-evolved” and populations evolved at ambient CO_2_ as “ambient-evolved”. All populations were grown in sterile microwell plates in 2mL of modified Tris-Acetate-Phosphate (TAP) media (Harris, 2009) at a standard temperature of 25 °C (Lindberg & Collins, 2020). The acetate was not added in this experiment so that CO_2_ was the only source of added carbon for the growing populations (TP media). Growing populations were maintained in mid‐log exponential growth by serial transfers of an inoculum (1% of each culture, ~10^4^ cells) to fresh medium three times per week. Details of TP media composition are in SI Table S1.

### 2.2 Experimental Design

To study the effect of environmental quality on the variation in trait values and combinations of trait values (hereafter referred to as multitrait phenotypes) expressed over all strains, we grew each of the single strain populations in a total of 8 different environments. For each environment, we defined environmental quality as the population growth rate that the particular environment allowed (Caruso, 2015; Caruso et al., 2016); the environments used here represented a range of qualities. For three weeks, populations were grown under low light (LL) conditions (30 ± 3 μmol photons m^−2^ s^−1^ (grow-lux lights), combined with either control (standard) media, media with reduced nutrients (Gorman & Levine, 1965), control media + high pCO_2_ (2000 ± 10 μatm CO_2_), or control media + high temperature (35°C ± 0.5 °C). We stress that the goal of this experiment was to study phenotypic variation under a wide range of growth rates in laboratory cultures – these do not, and are not intended to, represent or mimic natural environments, and buffered TP medium does not reproduce the carbonate chemistry of natural systems. This experiment was not designed to project expected trait combinations under carbon enrichment *per se*, but rather to give insight into how trait diversity itself is affected by environmental improvement vs. degradation relative to the recent evolutionary history of the populations. See Table 1 for environmental quality as measured by population growth rate for each environment. Standard pCO_2_ was 430 ± 10 μatm and standard temperature was 25 °C (± 0.5 °C); these temperatures and pCO_2_ conditions were applied unless otherwise noted. At the end of three weeks of acclimation to the environments (~ 9 growth cycles) we measured traits associated with the ecological function of aquatic primary producers: population growth rate (μ), gross photosynthesis rate (*GPR*), respiration rate (*R*), carbon use efficiency (CUE), cell size, relative chlorophyll autofluorescence per cell (*Chl*), intracellular reactive oxygen species content (ROS), and growth responses to the presence of conspecifics. Other than growth responses to conspecifics, these traits have previously been shown to vary over this populations in high vs. ambient CO_2_ environments in TAP media under high light at 25 °C (Lindberg & Collins, 2020). Our rationale for using multitrait phenotypes is that the ecological function of primary producers depends on several traits and their correlations – a greater range of multitrait phenotypes can indicate more strategies for responding to environmental change. The range of multitrait phenotypes expressed may be important in terms of projecting how trophic or other interactions may be affected in aquatic systems. To increase the range of environmental qualities in this study, we carried out a second experiment where the cells were acclimated for three weeks (~ 9 growth cycles) to the same 4 environmental conditions, this time under high light (HL) conditions (100 ± 3 μmol photons m^−2^ s^−1^), with the same traits measured after 3 weeks (Figure 2).

**Figure 2.**
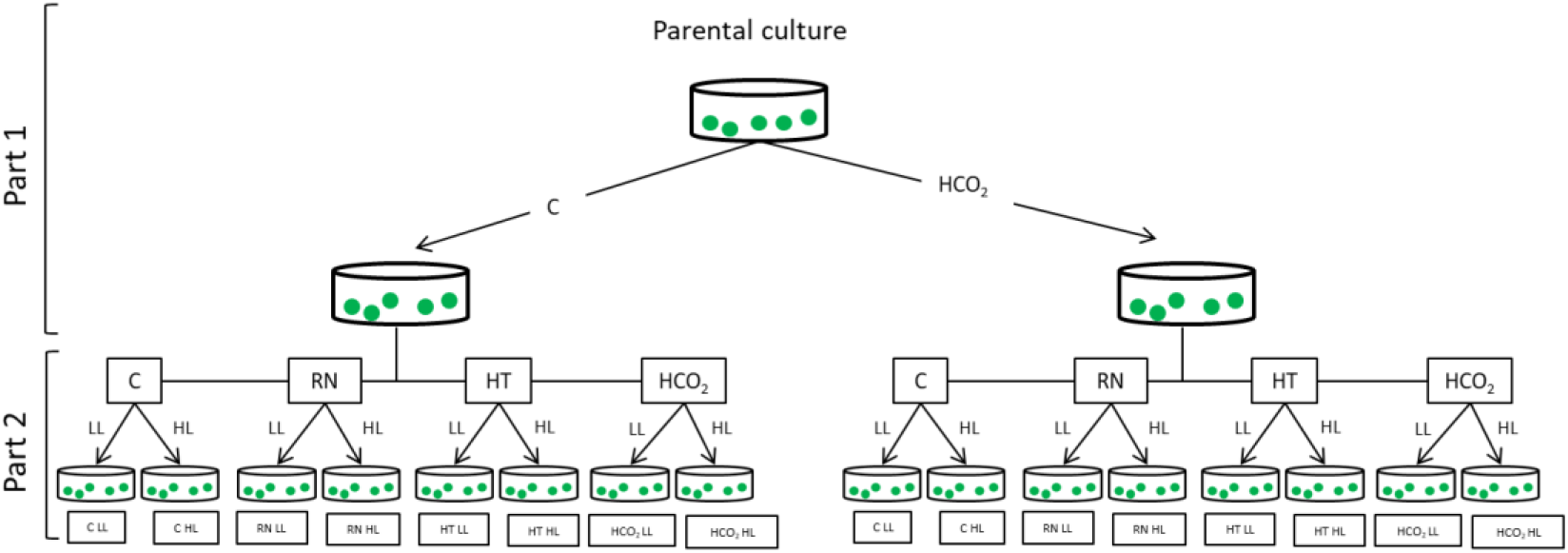
Schematic representation of experimental design. The experiment consists of two parts: In the first part (Part 1), the populations are exposed for 90 growth cycles to two different levels of CO_2_: ambient CO_2_ (430 μatm. CO_2_) and high CO_2_ (2000 μatm CO_2_) (Lindberg & Collins, 2020). Part 1 was carried out before the experiments in this study. Only a single strain is shown here for simplicity; the above was repeated for 3 strains, with 3 independent populations per strain per CO_2_ history. In the second part (this study), populations were grown for three weeks at low light (LL) or high light (HL) under the following environmental conditions (see Table 2): Control, Reduced nutrients (RN), High temperature (HT) and High CO_2_ (HCO_2_). Traits measurements and co-culture experiments were conducted at the end of the three week period. (See Figure 3 for co-culture details).

**Table 1.**
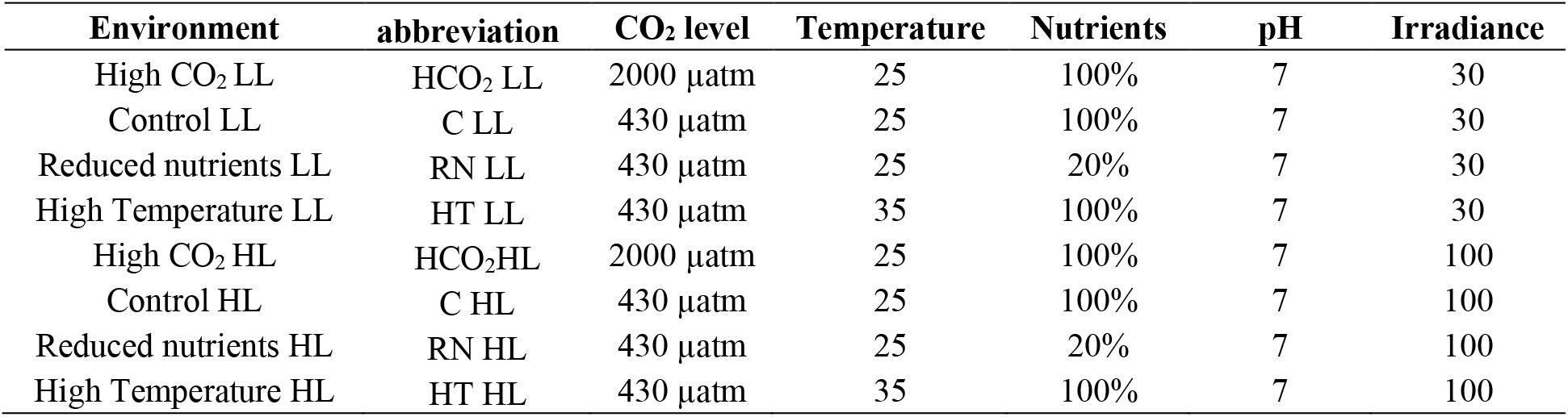
Details of environments used in the experiment. Units: CO_2_ level (μatm); Temperature (°C); Nutrients (percentage with respect to the control); LL/HL (continuous photon flux μmol photons m^−2^ s^−1^).

Our main goal in this study is assess how environmental quality affects first, variation in multitrait phenotypes and second, growth in the presence of conspecifics, in a model microalga (*C. reinhardtii).* To disentangle this from trait variation attributable to strain and recent evolutionary history, we include several strains and two known evolutionary histories in this experiment.

**Table 2.** Summary of effect sizes and significance levels of three-way ANOVAS for effects of genotype, experimental environment and previous history on growth rate (μ), gross photosynthesis rate (GPR), respiration (R), size, relative chlorophyll autofluorescence per cell (*Chl*) reactive oxygen species production (ROS) and carbon use efficiency (CUE). Asterisks represent significance level: < 0.001 ‘***’; < 0.001 ‘**’; < 0.01; ‘*’< 0.05. All data are in SI Table S2.

| Source of variation | Phenotypic variables |  |  |  |  |  |  |
| --- | --- | --- | --- | --- | --- | --- | --- |
| | $\mu$ | GPR | R | Size | *Chl* | ROS | CUE |
| Genotype | 0.014** | 0.119*** | 0.083*** | 0.010<sup>ns</sup> | 0.007<sup>ns</sup> | 0.032** | 0.014<sup>ns</sup> |
| Experimental environment | 0.727*** | 0.308*** | 0.203*** | 0.256*** | 0.484*** | 0.132*** | 0.151** |
| Previous History | 0.007* | 0.002<sup>ns</sup> | 0.001<sup>ns</sup> | 0.055*** | 0.000<sup>ns</sup> | 0.003 | 0.002<sup>ns</sup> |
| Genotype x Experimental environment | 0.101*** | 0.133*** | 0.139* | 0.275*** | 0.134*** | 0.389*** | 0.134<sup>ns</sup> |
| Genotype x Previous History | 0.003<sup>ns</sup> | 0.000<sup>ns</sup> | 0.000<sup>ns</sup> | 0.001<sup>ns</sup> | 0.008<sup>ns</sup> | 0.032*** | 0.002<sup>ns</sup> |
| Experimental environment x Previous History | 0.009<sup>ns</sup> | 0.066** | 0.051<sup>ns</sup> | 0.086*** | 0.036<sup>ns</sup> | 0.149*** | 0.008<sup>ns</sup> |
| Genotype x Experimental environment x Previous History | 0.019<sup>ns</sup> | 0.085* | 0.104<sup>ns</sup> | 0.045<sup>ns</sup> | 0.088** | 0.277*** | 0.105<sup>ns</sup> |

### 2.3 Population growth rate

The population growth rate (μ) of *C. reindhardtii* was estimated in log-phase, using the equation:

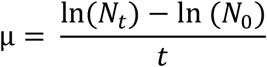

with *N*_t_ number of cells after a time period *t*, *N*_0_ number of cells at inoculation, and *t* the time passed in days. Cell numbers were determined using a FACSCanto II (BD Biosciences) flow cytometer calibrated with Cytometer Setup and Tracking (CS&T) beads and the data were acquired with the BD FACSDiva v6 software (Brennan & Collins, 2015). Each culture was counted at least twice. We made standard curves using flow cytometry cell counts and a spectrophotometer (Spark Control Tecan ^(R)^) in order to measure lineage frequencies during co-culture experiments. We detected the excitation/emission wavelength of 455/680 nm of the culture using a plate reader and fit a linear regression: CN= ((F_455-680_) / 0.027) - 4820 (*r*^2^= 0.868, *n*= 30), where F_455-680_ is the excitation/emission wavelength of 455/680 nm of the cell culture in a spectrophotometer (Spark Control Tecan ^(R)^).

### 2.4 Estimation of cell size and relative chlorophyll autofluorescence

Cell size was measured using flow cytometry FACSCanto II (BD Biosciences). Forward scatter area was related to cell diameter by a standard curve using microbeads. The diameter was calculated as μM = x 9.01×10^−5^ + 0.28873 where x is the value for forward scatter. The average cell diameter per population was based on a sample of at least 10000 cells. Volume was estimated assuming cells are spheres.

Relative chlorophyll autofluorescence per cell (*Chl*) was measured using flow cytometry FACSCanto II (BD Biosciences). Specifically, the relative chlorophyll autofluorescence intensity was detected in the PerCP-Cy5.5 channel (Ex-Max 488 nm/Em-Max long pass (LP) 670 – 725 nm). Samples were run from 96 well plates, at flow rates of 1μl/second.

### 2.5 Photosynthesis and respiration measurements

Gross photosynthetic rate (*GPR*) was calculated from oxygen production using an SDR SensorDish^®^ Reader. Samples (2.5 mL per assay) were initially incubated for 20 min in darkness at the relevant temperature and CO_2_ levels, then exposed for 5 min to 30 ± 3 μmol photons m^−2^ s^−1^ (LL) or 100 ± 3 μmol photons m^−2^ s^−1^ (HL). Respiration rate (*R*) was measured as O_2_ consumption over five minutes in the dark immediately following the photosynthesis measurements. All SDR measurements were made in the same incubators under the same conditions as the populations had acclimated to for three weeks prior to the trait measurements.

Carbon use efficiency (*CUE*) was calculated as:

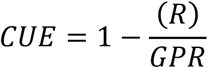

### 2.6 Reactive oxygen species assay

Relative intracellular reactive oxygen species (ROS) levels were determined using the dye 2,7-dichlorofluorescein diacetate (H_2_DCFDA) solubilized in ethanol (Gomes et al., 2017; Stoiber et al., 2013; Szivák et al., 2009). The DCFH-DA probe is membrane permeable, and has been shown to be reactive to a variety of ROS, particularly H_2_O_2_ and peroxide-derived oxidants (Imrich et al., 1999), but nonreactive to superoxide radicals (Lebel et al., 1992). We inoculated ~10^6^ cells per well from the cultures exposed to different experimental environments and the samples were incubated in the dark for 5 h after the addition of H_2_DCFDA. The final concentration in each well of dye was 2.5 μM H_2_DCFDA. This concentration of dye is less than used in previously studies with microalgae (Gomes et al., 2017; Knauert & Knauer, 2008; Stoiber et al., 2013; Szivák et al., 2009), but resulted in a clear signal using our microplate reader. The resulting ROS product was quantified by fluorescence using a microplate reader (Spark Control Tecan ^(R)^) at an excitation/emission wavelength of 488/525 nm (Gomes et al., 2017; Szivák et al., 2009).

### 2.7 Growth response to conspecifics

To test how strains with different previous evolutionary histories responded to the presence of a closely-related conspecific in each environment, we used ThinCert™ cell culture inserts (Figure 3). An insert is a 0.4 μm-membrane that is suspended in the individual wells of a 12-well plate. The insert permits extracellular products including nutrients to diffuse but prevents cells from doing so. Here, we used three *C. reinhardtii* strains (each strain has independently evolved populations with different histories, having been either evolved at high or ambient pCO_2_). For all assays, we inoculated the compartments inside and outside the insert with the same number of cells (approximately 7500 cells). For the mono-culture ‘control’ conditions, both compartments were inoculated with the same population. For the co-culture assays, the populations were grown in a full factorial setting, with three (low light conditions) or four (high light conditions) biological replicates (each biological replicate is an independently-evolved population of a given strain) and three technical replicates (the same population or set of populations grown in three independent wells). The use of the same strain with different evolutionary CO_2_ history in the experiment gives us information about the effect of selection history on the ability to respond to the presence of a closely related conspecific. Samples were distributed so that no one lineage was present solely in the outside or inside compartment in either combination. Growth rates were checked for each compartment separately. Under these conditions, resource competition is virtually absent, so that differences in growth in the presence of self vs non-self on the other side of the membrane indicate a plastic adjustment of population growth rate in response to the identity of the potential competitor. Growth rate response was calculated as:

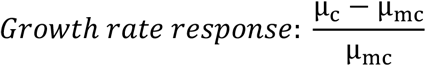

**Figure 3.**
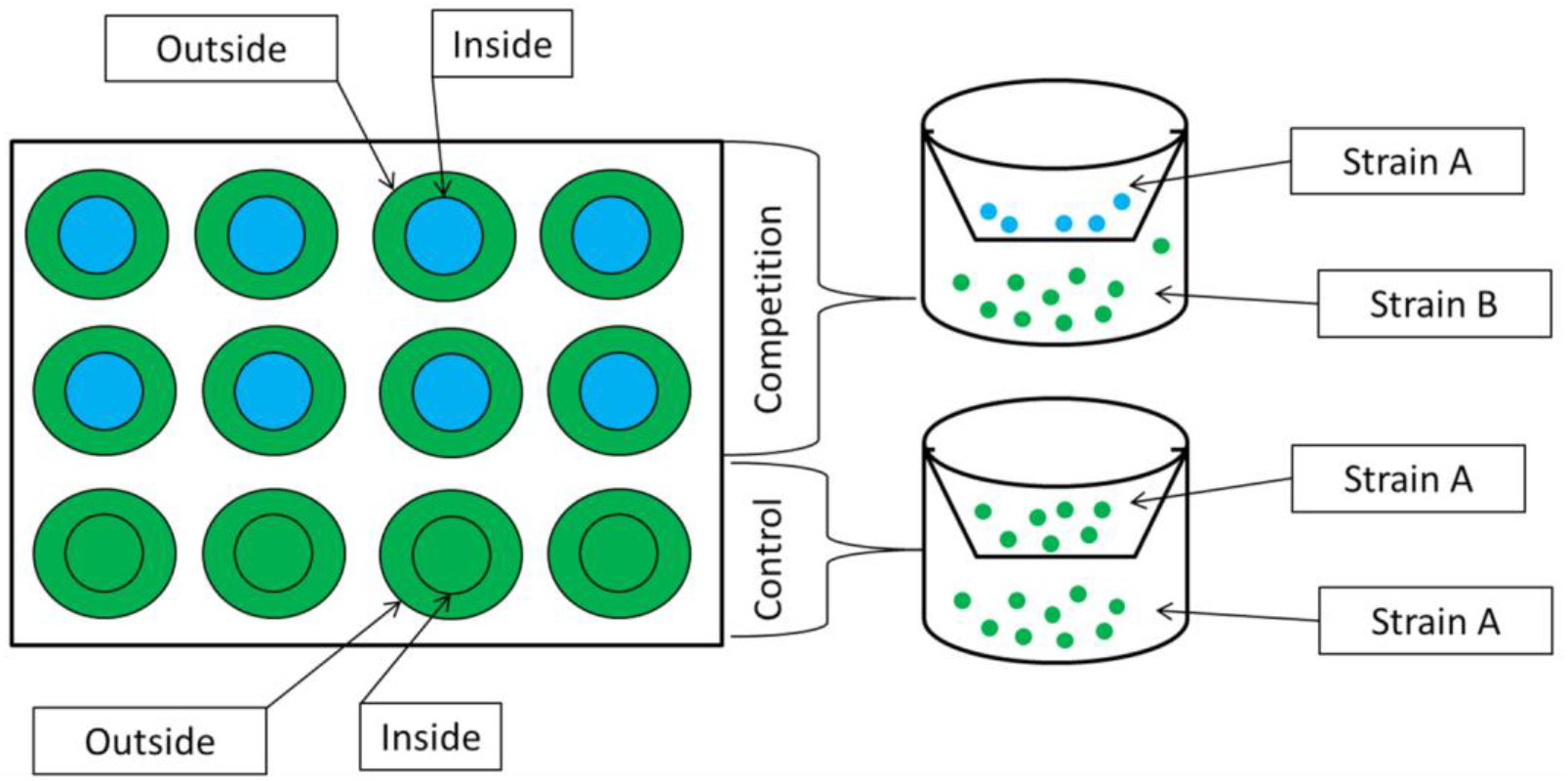
Co-culture experiment in microwell plates using ThinCert membranes. Samples were cultured in wells divided by a semi-permeable membrane (ThinCert). Control experiments were inoculated with the same strain inside/outside the membrane. Competition experiments were inoculated with cells of the same strain with different evolutionary histories (high-evolved or ambient-evolved).

Where μc is the population growth rate of a population when is growing in the presence of non-self and μmc is the growth rate of mono-culture ‘control’ conditions (the same population is growing inside and outside of ThinCert™).

### 2.8 Statistical analysis

For the monocultures, the objective of this study was to understand how variation in trait values (μ, *GPR*, *R*, cell size, ROS production, CUE and *Chl*) depends on environmental quality, strain and previous evolutionary history. To this, a three-way ANOVA was performed (model: y = overall mean + strain + experimental environment + previous CO_2_ history + strain × experimental environment + strain × previous CO_2_ history + experimental environment × previous CO_2_ history + strain × experimental environment × previous CO_2_ history + error). The factor “strain” corresponds to the three strains used in the experiment, the factor “experimental environment” corresponds with the eight combination of the ambient and irradiance tested in the experiment, and finally, previous CO_2_ history corresponds to whether the population evolved at high or ambient CO_2_ prior to this experiment. A similar analysis was performed for other traits: GPR, R, CUE, *Chl*, cell size, and ROS.

We used principle components analysis (PCA) to understand how trait values were related to each other to form multitrait phenotypes in each of our 8 environments. The traits included in the PCA were: GPR, R, CUE, *Chl*, cell size, and ROS. To allow us to investigate the role of evolutionary histories, a single PCA was performed with all populations, so that all populations were represented in a single PCA space. The analysis was performed in R Core Team (2013) using the FactoMineR package.

To test whether the position of a focal population relative to the ThinCert™ cell culture inserts affected growth responses to conspecifics during indirect co-culture, we used a four-way ANOVA with the following explanatory factors: position, strain, previous CO_2_ history and experimental environment. Here, “position” corresponds to the position of the population relative to the ThinCert™ membrane (inside/outside).

Finally, to explore how the growth responses to conspecifics were related to the extent of multitrait space available, as well as by their growth rates in monoculture, in a given environment, we compared models of growth response to conspecifics with and without interaction terms between monoculture growth rate and PCA ellipse volume using the Akaike information criterion with a correction for small sample sizes (AICc; Cavanaugh, 1997) using dredge tool from the package “MuMIn” in R (R Core Team, 2013).

## 3. RESULTS

To ensure that the environments used in this study represented a range of qualities for *C. reinhardtii*, we measured population growth rate in 8 different environments for 3 different strains, where each strain had populations with evolutionary histories of both high CO_2_ or ambient CO_2_ (Figure 2). We found that population growth rate depended on environment (SI Table S2, *df* = 7 and 96, *F* = 82.00, *p*<0.001), demonstrating that the environments in this study represented a range of qualities for these *C. reinhardtii* populations. Environment-specific growth rates depended on strain (SI Table S2, *df* = 2 and 96, *F* = 5.58, *p*<0.05) and previous CO_2_ history (SI Table S2, *df* = 1 and 96, *F* = 5.23, *p*< 0.05) (Figure 4, Table 2, SI Table S2). Experimental environment and the interaction between experimental environment and strain explained most of the population growth rate, with evolutionary history having a smaller effect size (Table 2). Here, the highest values of μ were found when the cells were grown under high light and high CO_2_, whereas the lowest values of μ occurred under low light and high temperature (Figure 4). This ranking of environmental qualities is broadly consistent with previous experiments with *C. reinhardtii* under laboratory conditions (Brennan & Collins, 2015), and with other phytoplankton (e.g. Thomas *et al.*, 2017).

**Figure 4.**
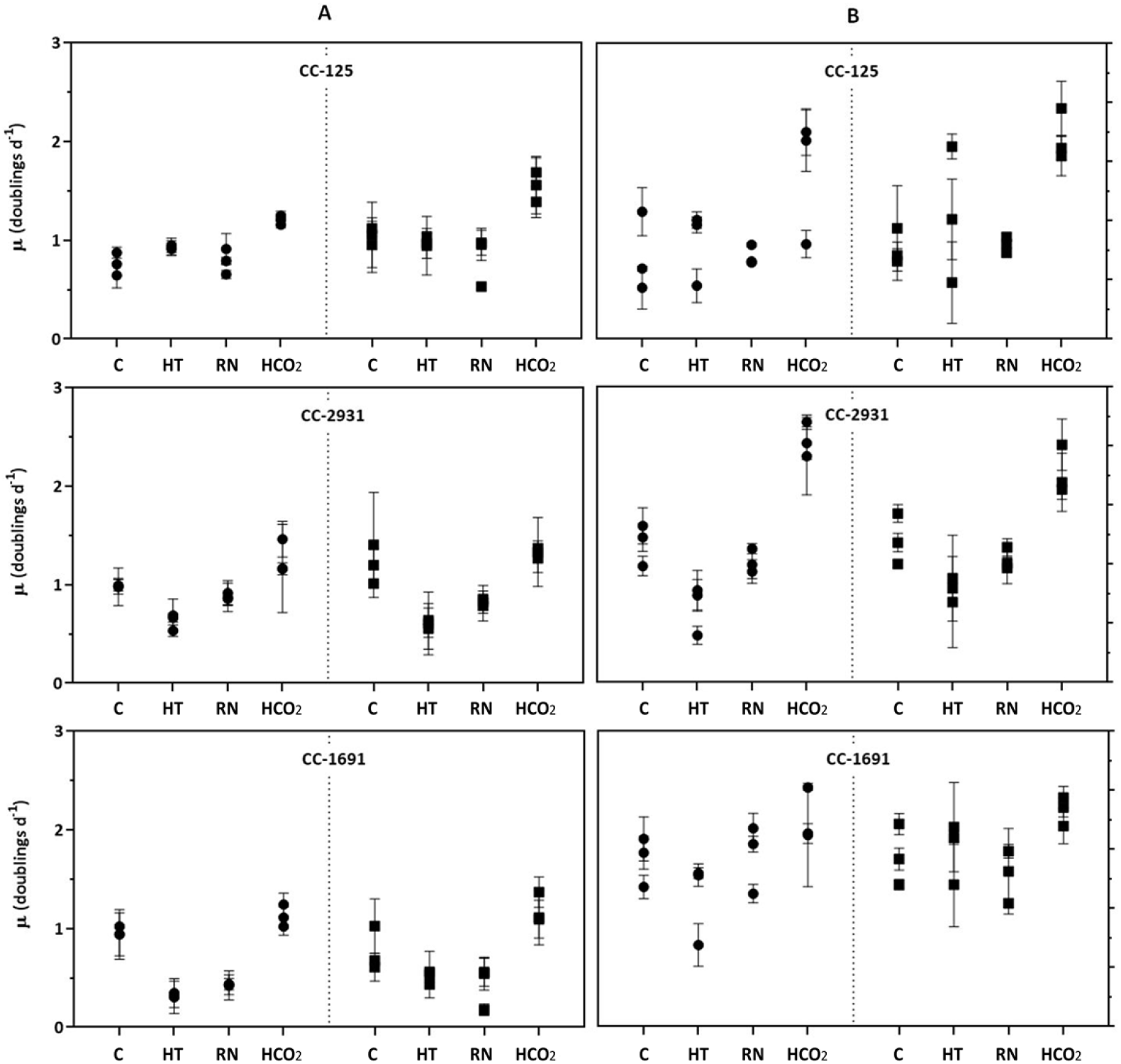
Population growth rates of *C. reinhardtii* in each experimental environment (C: control; HT: High temperature; RN: Reduced nutrients; HCO_2_: High CO_2_) in the LL (Column A) and HL (Column B). The symbols represent evolutionary history: ambient-CO_2_-evolved populations (circles) and high-CO_2_-evolved populations (squares). Each symbol represents an independently-evolved replicate population of a given strain, with three independent replicate measurements per independently-evolved population (mean ± SD, *n* = 3).

To understand how traits other than population growth were affected by environmental quality, and how this varied by strain and evolutionary history, we measured values for the following traits: gross photosynthesis rate, respiration rate, cell size, relative chlorophyll autofluorescence per cell, and intracellular reactive oxygen. We also calculated carbon use efficiency (CUE), which represents the fraction of carbon available for growth once the catabolic demands of the cell have been met (Raven & Geider, 1988), and which can be affected by temperature and pCO_2_ in single celled photoautotrophs (García-Carreras et al., 2018; Lindberg & Collins, 2020) .These traits differed by strain and evolutionary history for these populations in the evolution experiment preceding this study (Lindberg & Collins, 2020). Over all traits, the most important factor for explaining trait values is consistently the environment they are grown in (Table 2, SI Table S2). The interaction between strain and environment explains some of the variation of traits other than CUE. Previous CO_2_ history and its interaction with strain have the least effect on trait values, except for ROS. This is consistent with previous studies on these strains under high and ambient pCO_2_ (Lindberg & Collins, 2020), where the only consistent response to growing in a high CO_2_ environment for hundreds of generations over all strains is an increase in ROS tolerance and related gene expression, and other traits evolve differently between strains but similarly within strains.

To understand how variation in multitrait phenotypes is related to environmental quality (Figure 1), we used principal components analysis (PCA) to quantify the multitrait space occupied over all strains and in each environment. 77% of the overall variance is explained by three principal components (PCs). The PCs are a combination of *GPR*, *R* and *CUE* (PC 1, 40.71%), size and *Chl* for PC 2 (18.91%), and finally, ROS for PC3 (17.54%) (SI Figure S1). Since the high CO_2_ environments represent either same-quality or ameliorated environments depending on evolutionary history, we consider the two evolutionary histories separately in a common PCA space. To quantify the multitrait space used in each environment for each evolutionary history, we use ellipse volume (95% confidence) that includes all strains with a given evolutionary history for each environment (Figure 5). We found that PCA ellipse volume increases with a growth rate for ambient-evolved populations (Figure 6), but there is no relationship between the PCA ellipse volume and growth rate for the high-evolved populations (Figure 6). This may be because, for the high-evolved populations, all environments are either equivalent to or worse than the one they evolved in, so they experience a relatively narrow range of qualities. Since both the high and ambient-evolved populations were from an experiment that intentionally used a high CO_2_ environment that maximized growth rate, we were unable to find an ameliorated environment for the high-evolved populations for the current study. In contrast, for the ambient-evolved populations, this experiment contains environments that are both substantially better and worse than the one they evolved in. The ambient-evolved populations also have a larger range (67.3) and coefficient of variation (137.6) of values over all for PCA ellipse volumes than do the high-evolved populations (range 19.9, CV 62.2), so that any relationship present is easier to detect in the ambient evolved lines.

**Figure 5.**
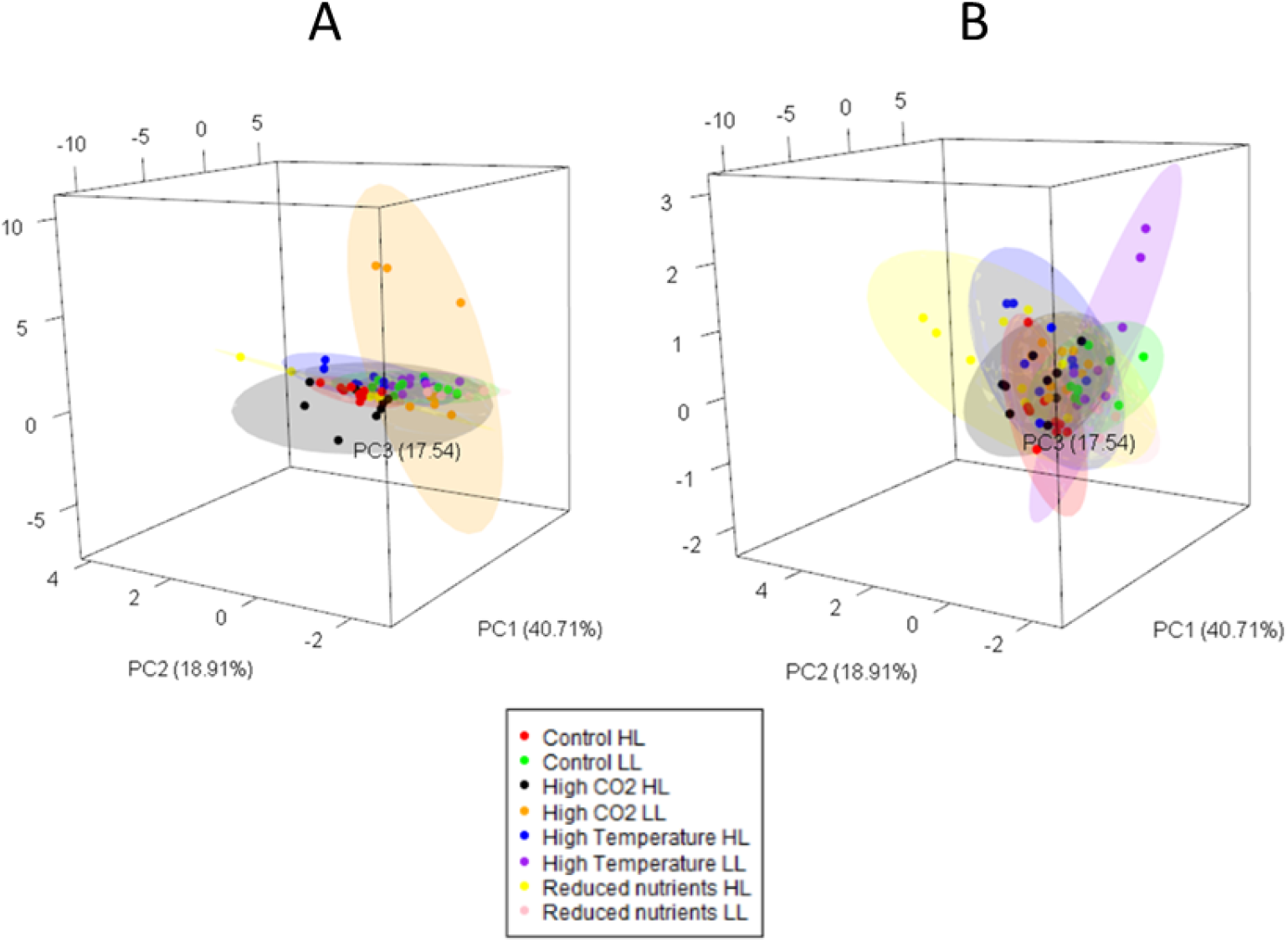
Principal Components Analysis (PCA) of the multitrait phenotypes of strains under different environmental conditions (A: ambient-evolved populations, B: high-evolved populations; all strains). Note that all strains were analysed together and projected into the same trait space; here we show duplicate identical trait spaces, each with strains of a single evolutionary history, for the purposes of visualization. Analysis based on the trait values for: *GPR*, *R*, *CUE*, size, *Chl* and *ROS*.

**Figure 6.**
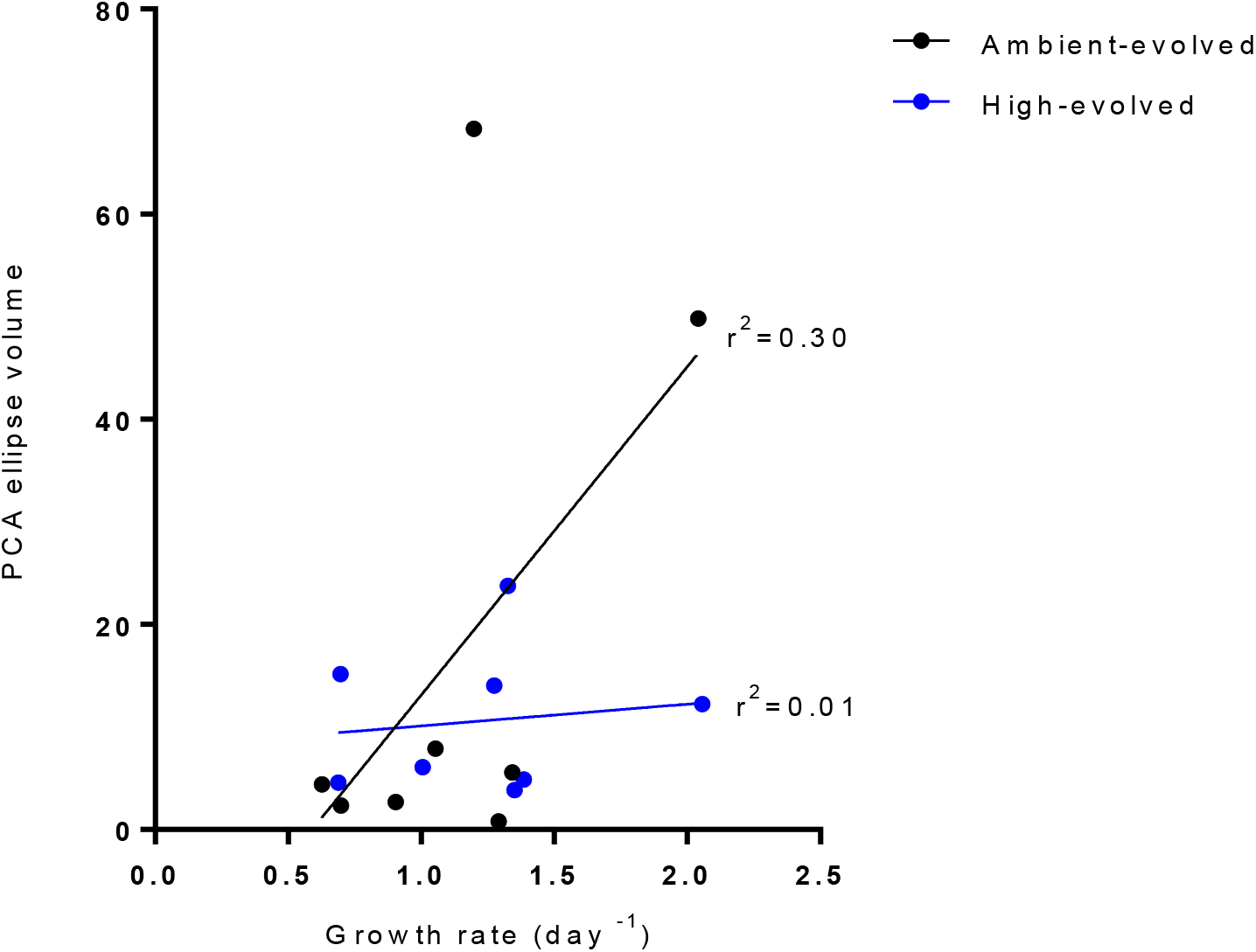
Relationship between PCA ellipse volume and population growth rate. PCA ellipse volume corresponds to the space that contains all populations of a common evolutionary history in each environment (3 populations for each of 3 strains) with 95% confidence.

To investigate the relationship between trait variation, environmental quality and responses to social cues, we test the hypothesis that there is more variation in growth responses to non-self conspecifics in higher-quality environments than in lower-quality ones. The reasoning behind this hypothesis is that during the long evolutionary history of the organisms prior to being brought into the laboratory, strategies for “winning” a competition could have included allocating energy away from population growth and towards other functions, such as nutrient storage, sex, or spore formation (Ghoul & Mitri, 2016; Hibbing et al., 2010). In contrast, in batch cultures that are transferred frequently, overgrowing competitors is the only strategy for winning competitions (Collins et al., 2019). We reason that in lower-quality environments, fewer energy allocation strategies can be used, while in higher-quality environments, more energy-allocation strategies exist, which is consistent with the positive relationship detected in this study between environmental quality and PCA ellipse volume in the ambient-evolved populations. We used indirect co-culture experiments to test whether the expected relationship between responses to the presence of closely-related non-self populations and the amount of multitrait phenotype space used could be detected.

High- and ambient-evolved populations of the same strain were grown in indirect co-culture in the 8 environments, and growth in the presence of conspecifics was compared to growth in the presence of self only for each population. Position relative to the ThinCert™ (inside vs. outside of membrane) does not significantly affect population growth rate (SI Table S3). Environment is the main determinant of growth (*df* = 7 and 114, *F* = 9.46, *p*< 0.01). The lineage frequencies in co-culture can be affected by two factors in this experiment, one of which corresponds to the growth rate of a focal population when grown in the presence of self only in that environment, and which reflects the growth rate due to abiotic environmental cues alone, such as different temperatures and nutrient levels. The second factor uses the range of phenotypes expressed over all populations with a shared evolutionary history in that environment, as an indication of the range of possible phenotypes in a given environment given very similar genetic capabilities. First, and as expected, the growth rate of a population in monoculture in that environment affects the frequency of that population in indirect co-culture, with faster-growing populations often, but not always, reaching higher frequencies than slower-growing ones. However, 40% of populations that reached higher frequencies in subdivided wells actually had *lower* monoculture growth rates (growth rate in the presence of self) than the population on the other side of the division (Table 3). This is because growth in the presence of a non-self conspecific is sometimes different than growth in the presence of a self population in indirect co-culture (SI Figure S2). Here, cell densities are initially low, and population growth rates are calculated well before cultures approach carrying capacity, so we suggest that this indicates that *C.reinhardtii* has the capacity to change population growth rate in responses to social cues from conspecifics. The PCA ellipse volume in this study represents the multitrait phenotypes available over all populations with a given CO_2_ evolution history in that environment. Here, for ambient-evolved populations, the larger the PCA ellipse volume for an environment and evolutionary history, the less the growth rate of a population in monoculture predicts its growth response to conspecifics in that environment. This could be because ambient-evolved populations have more trait variation than high-evolved populations overall, which is perhaps due to only ambient-evolved populations experiencing novel environmental improvement during this study. Variation in μc can be explained as:

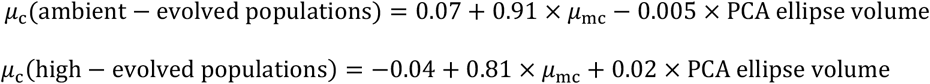

**Table 3.**
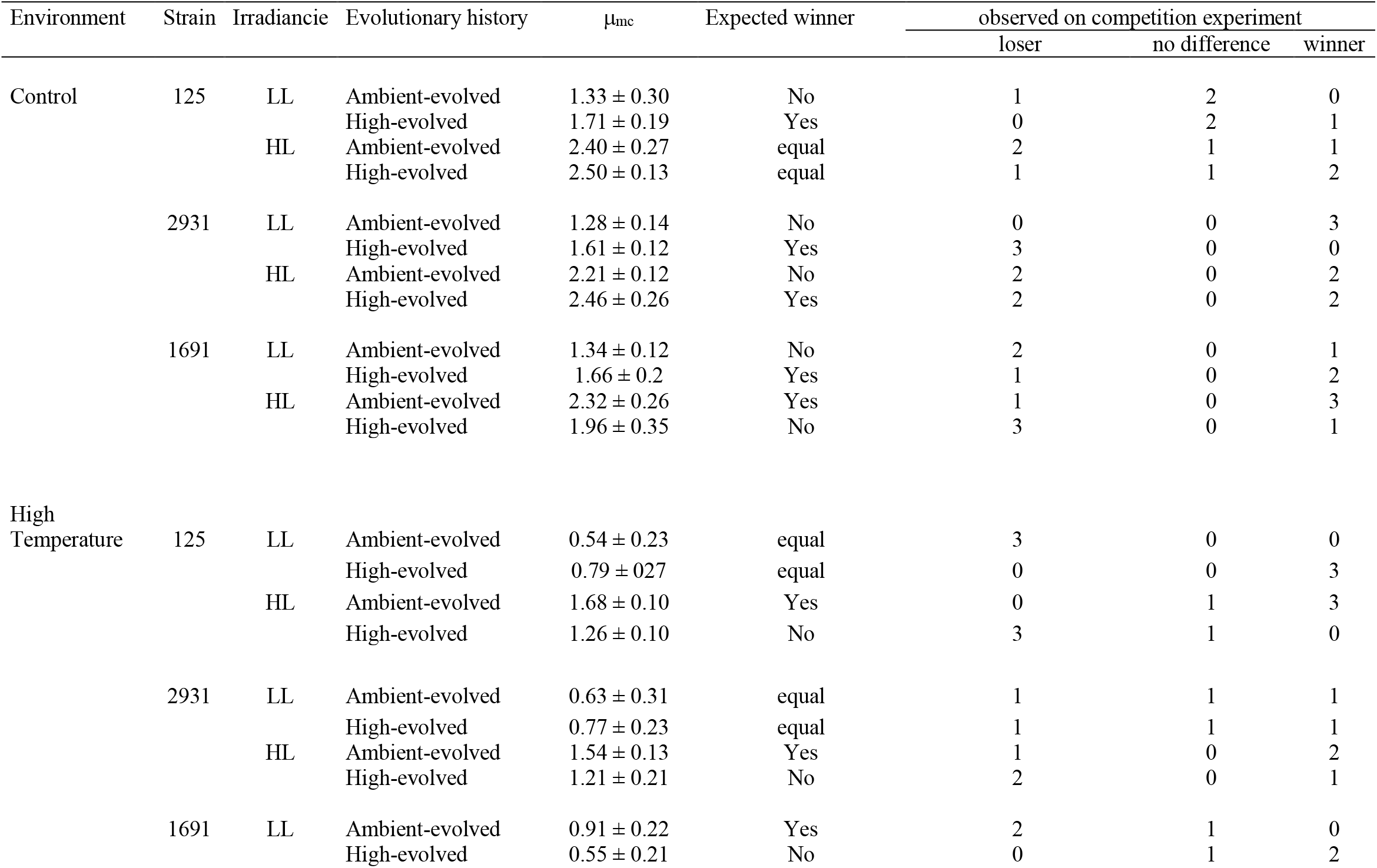
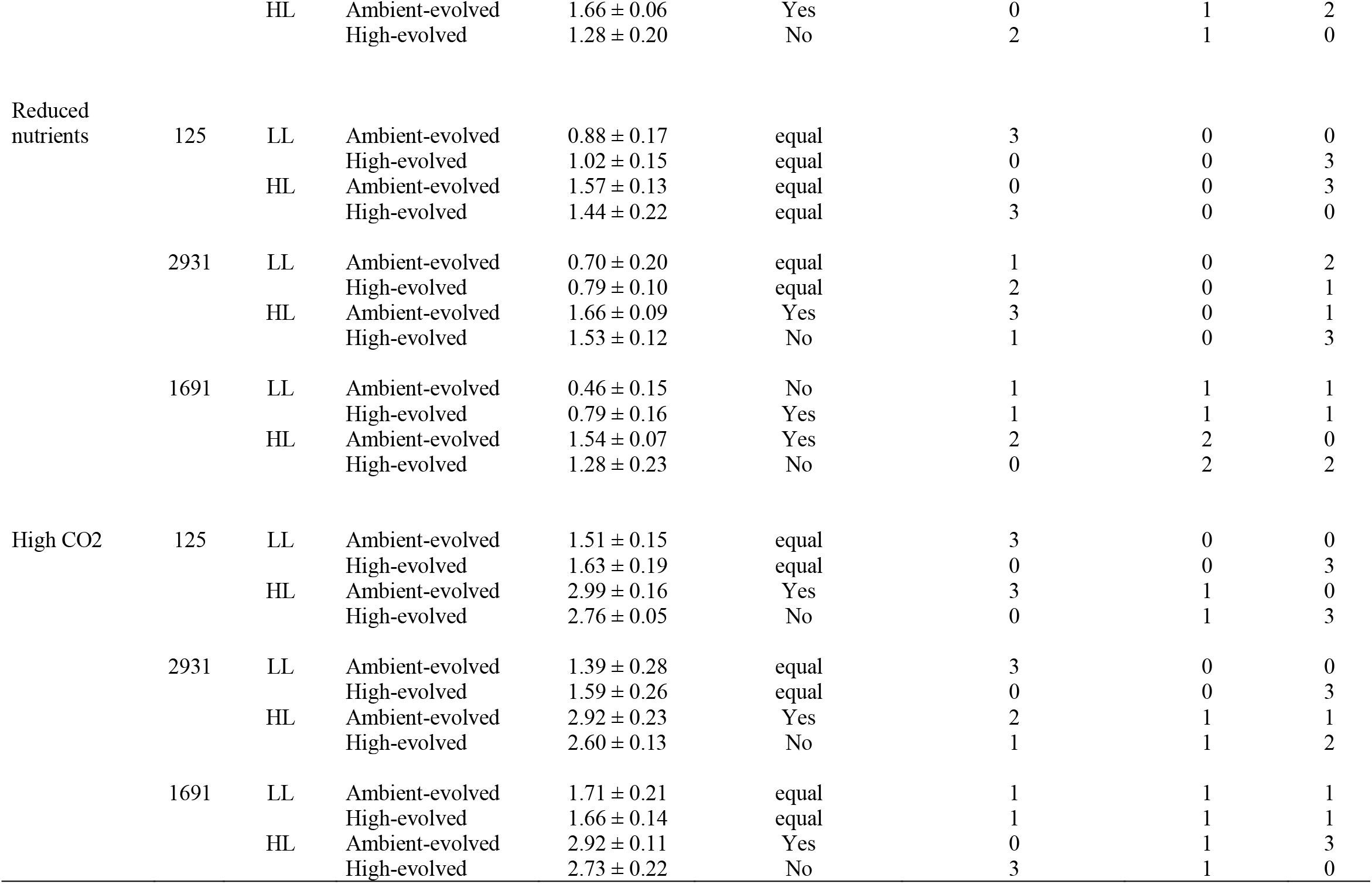
Results of indirect co-culture experiments. Comparison between expected winners (higher frequency strain based on growth in monoculture) and observed winner in competition experiment (higher frequency strain observed in co-culture) (see SI Figure S2).

The breakdown in predictability of lineage frequency from monoculture growth plays out mainly in the high CO_2_ environment, where high- and ambient-evolved populations of the same strain often have similar population growth rates in monoculture. Here, the expectation is that they would have equal abilities to respond to the presence of non-self conspecifics. However, the high-evolved populations generally have higher frequencies than expected in co-culture (Table 3), which can be explained by the ambient-evolved populations sometimes lowering their growth rates in the presence of non-self cues. This is in contrast with previous experiments with high CO_2_ evolved *C.reinhardtii*, which had lower than expected competitive abilities (Collins, 2010) though previous experiments only examined two environments, and did not use high CO_2_ conditions that maximized cell division rates for hundreds of generations. The results here are consistent with a recent study in a marine picoplanker (Collins & Schaum, 2019), where there was more variation in responses to non-self conspecifics in high CO_2_ (higher-quality) environments than in ambient CO_2_ (lower quality) environments.

## 4. DISCUSSION

The main goal of this study was to link environmental quality with intraspecific trait variation. First, we found that for one evolutionary history (ambient-evolved populations), more phenotypic variation is expressed in better quality environments, in line with scenarios A and B in Figure 1. In contrast, high-evolved populations showed no such trend. We tentatively ascribe this to the multitrait variation being greater in the ambient-evolved populations overall, or to high pCO_2_ environments being novel high quality environments for these populations, or a combination of the two. Subsequently, we explored how environmental quality and phenotypic variation affected lineage frequencies in fast-growing indirect co-culture. We found that when populations are in high-quality environments, monoculture growth rate becomes a less reliable predictor of lineage frequencies than it is in poor quality or non-novel environments.

### 4.1 Environmental quality affects phenotypic variation

We hypothesized that shifts in environmental quality can affect the range of phenotypes expressed in a group of closely-related populations (Figure 1). In extremely degraded or initially toxic environments, we expect a low number of strategies (low phenotypic variation), as few phenotypic solutions allow non-negative population growth rates. Plastic or evolutionary responses to these environments, when studied, often involve increasing tolerance to a toxin or stress (Andersson, 2006; Andersson & Levin, 1999; Coustau et al., 2000). In contrast, ameliorated environments allow higher population growth rates, and evolved values for many key traits differ widely for populations with similar growth rates (Lindberg & Collins 2020). This suggests that a wider range of multitrait phenotypes are expressed when two conditions are met: first, when there are more resources (higher nutrients and more light, for example), and second, when the increase in resources is novel for the populations (elevated CO_2_ for the ambient evolved populations, for example). For example, lower population growth rates and higher reactive oxygen tolerance can evolve in CO_2_-enriched environments to avoid the accumulation of cellular damage (Collins, 2016; Lindberg & Collins, 2020; Litchman et al., 2015; Low-Décarie et al., 2011; O’Donnell et al., 2018; Schaum & Collins, 2014), but other traits, such as cell size and photosynthesis rates, vary between populations that have adapted to the same environments (Lindberg & Collins, 2020). In line with this, we observed that the growth environments used in this experiment explained most of the changes in trait values that we measured. For the ambient-evolved populations, we observed a positive relationship between the number of strategies (estimated by PCA ellipse volume) expressed over all genotypes in a given environment, and the quality of that environment. These populations experienced both novel poor and novel high-quality environments during this study. Environments that are qualitatively different, but of similar overall quality, don’t increase overall phenotypic variation, even though they do change which phenotypes are more fit. In contrast, environmental amelioration increases phenotypic variation and also affects which phenotypes are most fit. The positive relationship in the ambient-evolved populations is largely driven by the extremely large ellipse volumes in the two high pCO_2_, environments, which are ameliorated environments for these populations. The ambient evolved populations have much higher growth rates in the high CO_2_ environments than they do in the corresponding low CO_2_ environments at a given light level (Low light: *df* = 52, *t* = 6.03, *p*< 0.01; High light: *df* = 52, *t* = 6.78, *p*< 0.01). While the high-evolved populations show similar differences in growth rates between the control and high CO_2_ environments for each light level, high pCO_2_ is not an ameliorated environment for these populations, as they had previously evolved under high pCO_2_.

Taken together, these data suggest that ameliorated environments (such as CO_2_ enrichment, temperature increase in some parts of the world, or nutrient enrichment) may at least initially increase variation more than expected based on increases to system carrying capacity (extra energy input) alone (Biswas et al., 2017; Burson et al., 2018; Gudmundsdottir et al., 2011; Richardson et al., 2019; Stevenson et al., 2008).

### 4.2 Phenotypic variation and responses to conspecifics

In this experiment, we found that monoculture growth rate is a poor predictor of lineage frequencies during rapid growth in high-quality environments. This has been previously observed in several studies of marine phytoplankton (Tatters *et al.*, 2013; Schaum & Collins, 2014; Wolf *et al.*, 2019). Using monoculture growth to predict lineage frequencies in the absence of resource competition has two complications that can come into play: first, if strains modulate their growth rate in response to cues from non-self organisms, then growth in monoculture may not predict growth in mixed culture. Second, cell division rates are only one component of fitness in the natural world, and few (if any) organisms evolved in environments where cell division rates were the only trait underlying fitness. Indeed, microbes often slow or arrest growth (Del Giorgio & Gasol, 2008; David L Kirchman, 2016; Smriga et al., 2014; Tada et al., 2010, 2013). Known cases where lowered population growth rates are adaptive include spore formation (Roszak & Colwell, 1987; Sussman & Douthit, 1973; Whittington et al., 2004) and predator or virus avoidance (Frickel et al., 2016; Lennon et al., 2007; Lürling, 1999; Ploug et al., 1999; Yoshida et al., 2004). It is thus unsurprising that in organisms evolved for countless generations under variable conditions before they were domesticated into laboratory model systems, that lineages may modulate their growth rates in response to cues other than resource availability. Similarly, we should expect that organisms evolved in natural communities prior to being isolated for laboratory studies can (still) detect and respond to the presence of non-self organisms.

The consequences of intraspecific phenotypic variability increasing with environmental amelioration could affect the potential for rapid evolution from standing variation (genotype sorting) within species experiencing rapid environmental shifts. Rapid environmental amelioration could substantially increase intraspecific trait variation during periods where populations are growing at their maximum rate, which could speed up evolution within species. In contrast, rapid environmental degradation has the potential to decrease intraspecific variation, thus slowing evolutionary responses (this remains to be tested, as we did not use any severely degraded environments in this study). In the case of environmental amelioration, this effect may be exacerbated by lineages also modulating their growth rate in response to social environment, which can decrease the predictability of evolution based on monoculture experiments. One important caveat is that our experiment did not use any environments where growth was severely slowed or which contained toxins, all of these environments were “good enough” for populations to have non-negative growth rates. We also did not include novel qualitative environmental challenges, such as different forms of nutrients. Our study used population densities where competition for resources were low, and co-cultured populations were growing at their maximum rate of increase given the environment. Because of this, these findings are especially relevant for understanding how environmental quality affects the potential for genotype sorting during rapid, density-independent population growth, such as during the early stages of phytoplankton blooms, or during laboratory experiments that propagate populations by batch culture, and then use competitive assays to quantify fitness gain.

We have found that environmental quality, measured as the maximum population growth rate that an environment allows, can affect trait variation and lineage frequencies in a model microalga under laboratory conditions. Should this translate into natural populations experiencing rapid environmental shifts, this relationship between intraspecific trait variation and changes in environmental quality can affect the potential for rapid adaption within species.

## Acknowledgements

AF-M and IJM-J were supported by the project CGL2017‐87314‐P (*Ministerio de Economía, Industria y Competitividad*, *España*), IJM-J was also supported by the University of Malaga (*Plan Propio de Investigación y Transferenci*a, *España*). SC was supported by a Royal Society University Research Fellowship (UK).

## Conflict of Interest

The authors declare no conflict of interest.

## Author contribution

IJM-J performed the experiments, analyzed the data, wrote the paper, prepared figures and tables, and reviewed drafts of the paper; AF-M reviewed drafts of the paper; SC conceived and designed the experiments, analyzed the data, contributed reagents/materials/analysis tools, wrote the paper and reviewed drafts of the paper.

## Data availability statement

All relevant data are within the paper and its Supporting Information files.

## Notes

### Competing Interest Statement

The authors have declared no competing interest.

