## Supplementary material for "The role of changes in environmental quality in multitrait plastic responses to environmental and social change in the model microalga *Chlamydomonas reinhardtii*": SI

**Content**:

### Tables:

**Supplementary Table 1|** Molar composition of culture media.

**Supplementary Table 2|** Three-way-ANOVAs of single-strain monocultures testing the effect of experimental environment, genotype (strain), previous $CO_2$ history and the interactions on μ, GPR, R, size, *Chl*, ROS and CUE of *Chlamydomonas reinhardtii*.

**Supplementary Table 3|** Four-way ANOVAS for the comparison of the experimental environment, genotype (strain), previous $CO_2$ history and position on the growth rates of control competition experiment ($\mu_{mc}$) under low light (LL) and high light (HL).

### Figures:

**Supplementary Figure 1|** Percentage the explanation of each phenotypic trait on the three principal components (PCs) derived from the PCA analysis. The three PCs explained the 77% of the overall variance. Specifically, PC 1 explained 40.71%, PC 2 explained 18.91%, and PC3 explained 17.54%.

**Supplementary Figure 2|** Growth in the presence of a non-self population in indirect competition ($\mu_c$) relative to growth in the presence of a self population in indirect competition ($\mu_{mc}$) for ambient-evolved (A) and high-evolved (B) populations.

**Table S1.** Molar composition of culture media.

| Major components (mM) | |
|---|---|
| $NH_4^+$ | 7.48 |
| $K^+$ | 1.94 |
| $Na^+$ | 0.27 |
| $Ca^{2+}$ | 0.34 |
| $Mg2^+$ | 0.41 |
| $Cl^-$ | 8.22 |
| $SO_4^{2-}$ | 0.51 |
| $PO_4^{3-}$ | 1.00 |
| Tris | 20.0 |
| **Trace Componentes (μM)** | |
| $Fe^{2+}$ | 17.9 |
| $Zn^{2+}$ | 76.5 |
| $Cu^{2+}$ | 6.3 |
| $Co^{2+}$ | 6.8 |
| $Mn^{2+}$ | 25.6 |
| $Mo^{6+}$ | 6.2 |
| $BO_3^{3-}$ | 184 |
| EDTA | 134 |

**Table S2.** Three-way-ANOVAs of single-strain monocultures testing the effect of experimental environment, genotype (strain), previous $CO_2$ history and the interactions on μ, GPR, R, size, *Chl*, ROS and CUE. Units are: μ (doublings·$d^{-1}$), GPR (nmol·$O_2$·$10^6$ $cel^{-1}$·$h^{-1}$), R (nmol·$O_2$·$10^6$ $cel^{-1}$·$h^{-1}$), size (μM), *Chl* (Relative chlorophyll autofluorescence·$cell^{-1}$), ROS [$\lambda_{488-525}$·$10^6$ $cel^{-1}$·$h^{-1}$].

| Variable | Soure of variation | df | SS | MS | *F* | *P* |
|---|---|---|---|---|---|---|
| μ | Genotype | 2 | 0.478 | 0.239 | 5.58 | 0.01 |
| | Experimental environment | 7 | 24.573 | 3.510 | 82.00 | 0.00 |
| | Previous $CO_2$ History | 1 | 0.224 | 0.224 | 5.23 | 0.02 |
| | Genotype x Experimental environment | 14 | 3.405 | 0.243 | 5.68 | 0.00 |
| | Genotype x Previous $CO_2$ History | 2 | 0.089 | 0.044 | 1.04 | 0.36 |
| | Experimental environment x Previous $CO_2$ History | 7 | 0.304 | 0.043 | 1.02 | 0.42 |
| | Genotype x Experimental environment x Previous $CO_2$ History | 14 | 0.625 | 0.045 | 1.04 | 0.42 |
| | Error | 96 | 4.110 | 0.043 | | |
| GPR | Genotype | 2 | 3.29E+08 | 1.64E+08 | 1.99E+04 | 0.00 |
| | Experimental environment | 7 | 8.49E+08 | 1.21E+08 | 1.47E+04 | 0.00 |
| | Previous $CO_2$ History | 1 | 5.38E+06 | 5.38E+06 | 6.52E-01 | 0.42 |
| | Genotype x Experimental environment | 14 | 3.66E+08 | 2.62E+07 | 3.17E+03 | 0.00 |
| | Genotype x Previous $CO_2$ History | 2 | 7.16E+05 | 3.58E+05 | 4.30E-02 | 0.96 |
| | Experimental environment x Previous $CO_2$ History | 7 | 1.81E+08 | 2.59E+07 | 3.14E+03 | 0.01 |
| | Genotype x Experimental environment x Previous $CO_2$ History | 14 | 2.35E+08 | 1.68E+07 | 2.04E+03 | 0.02 |
| | Error | 96 | 7.93E+08 | 8.26E+06 | | |
| R | Genotype | 2 | 8.29E+07 | 4.15E+07 | 9.45 | 0.00 |
| | Experimental environment | 7 | 2.03E+08 | 2.91E+07 | 6.62 | 0.00 |
| | Previous $CO_2$ History | 1 | 6.63E+05 | 6.63E+05 | 0.15 | 0.70 |
| | Genotype x Experimental environment | 14 | 1.39E+08 | 9.96E+05 | 2.27 | 0.01 |
| | Genotype x Previous $CO_2$ History | 2 | 1.22E+06 | 6.12E+04 | 0.01 | 0.99 |

| | | | | | | |
|---|---|---|---|---|---|---|
| | Experimental environment x Previous $CO_2$ History | 7 | 5.09E+07 | 7.27E+06 | 1.66 | 0.13 |
| | Genotype x Experimental environment x Previous $CO_2$ History | 14 | 1.04E+07 | 7.45E+06 | 1.70 | 0.07 |
| | Error | 96 | 4.21E+08 | | | |
| Size | Genotype | 2 | 7.42E+04 | 3.71E+04 | 1.79 | 0.17 |
| | Experimental environment | 7 | 1.86E+06 | 2.66E+05 | 12.86 | 0.00 |
| | Previous $CO_2$ History | 1 | 3.97E+05 | 3.97E+05 | 19.21 | 0.00 |
| | Genotype x Experimental environment | 14 | 2.00E+06 | 1.43E+05 | 6.90 | 0.00 |
| | Genotype x Previous $CO_2$ History | 2 | 6.56E+03 | 3.28E+03 | 0.16 | 0.85 |
| | Experimental environment x Previous $CO_2$ History | 7 | 6.27E+05 | 8.96E+04 | 4.33 | 0.00 |
| | Genotype x Experimental environment x Previous $CO_2$ History | 14 | 3.29E+05 | 2.35E+04 | 1.14 | 0.34 |
| | Error | 96 | 1.99E+06 | | | |
| *Chl* | Genotype | 2 | 0.037 | 0.018 | 1.390 | 0.254 |
| | Experimental environment | 7 | 2.545 | 0.364 | 27.330 | 0.000 |
| | Previous $CO_2$ History | 1 | 0.018 | 0.002 | 0.130 | 0.715 |
| | Genotype x Experimental environment | 14 | 0.704 | 0.050 | 3.780 | 0.000 |
| | Genotype x Previous $CO_2$ History | 2 | 0.040 | 0.020 | 1.500 | 0.229 |
| | Experimental environment x Previous $CO_2$ History | 7 | 0.191 | 0.027 | 2.050 | 0.056 |
| | Genotype x Experimental environment x Previous $CO_2$ History | 14 | 0.460 | 0.033 | 2.470 | 0.005 |
| | Error | 96 | 1.277 | 0.013 | | |
| ROS | Genotype | 2 | 9.73E+06 | 4.86E+06 | 17.583 | 0.00 |
| | Experimental environment | 7 | 4.08E+07 | 5.83E+06 | 21.065 | 0.00 |
| | Previous $CO_2$ History | 1 | 8.16E+05 | 8.16E+05 | 2.951 | 0.09 |
| | Genotype x Experimental environment | 14 | 8.93E+07 | 6.38E+06 | 23.044 | 0.00 |
| | Genotype x Previous $CO_2$ History | 2 | 9.83E+06 | 4.91E+06 | 17.758 | 0.00 |
| | Experimental environment x Previous $CO_2$ History | 7 | 4.61E+07 | 6.58E+06 | 23.795 | 0.00 |
| | Genotype x Experimental environment x Previous $CO_2$ History | 14 | 8.54E+07 | 6.10E+06 | 22.041 | 0.00 |
| | Error | 96 | 2.66E+07 | 2.77E+05 | | |
| CUE | Genotype | 2 | 0.032 | 0.01602 | 1.164 | 0.32 |
| | Experimental environment | 7 | 0.3423 | 0.0489 | 3.554 | 0.00 |
| | Previous $CO_2$ History | 1 | 0.0053 | 0.00525 | 0.382 | 0.54 |
| | Genotype x Experimental environment | 14 | 0.3035 | 0.02168 | 1.575 | 0.10 |
| | Genotype x Previous $CO_2$ History | 2 | 0.005 | 0.00251 | 0.183 | 0.83 |
| | Experimental environment x Previous $CO_2$ History | 7 | 0.0177 | 0.00253 | 0.184 | 0.99 |
| | Genotype x Experimental environment x Previous $CO_2$ History | 14 | 0.2367 | 0.01691 | 1.229 | 0.27 |
| | Error | 96 | 13.211 | 0.01376 | | |

**Table S3.** Four-way ANOVAS for the comparison of the experimental environment, genotype (strain), previous $CO_2$ history and position on the growth rates of control competition experiment ($\mu_{mc}$) under low light (LL) and high light (HL).

| | Low light | | | | |
|---|---|---|---|---|---|
| Source of variation | SS | *df* | MS | F | *p* |
| A:Experimental environment | 23.33 | 3 | 7.78 | 163.26 | 0.00 |
| B:Position | 0.13 | 1 | 0.13 | 2.72 | 0.10 |
| C:Previous $CO_2$ history | 0.92 | 1 | 0.92 | 19.23 | 0.00 |
| D:Genotype | 0.16 | 2 | 0.08 | 1.65 | 0.20 |
| Interactions | | | | | |
| AB | 0.08 | 3 | 0.03 | 0.53 | 0.66 |
| AC | 0.58 | 3 | 0.19 | 4.03 | 0.01 |
| AD | 0.78 | 6 | 0.13 | 2.75 | 0.02 |
| BC | 0.00 | 1 | 0.00 | 0.00 | 0.98 |
| BD | 0.01 | 2 | 0.01 | 0.14 | 0.87 |
| CD | 0.17 | 2 | 0.08 | 1.75 | 0.18 |
| ABC | 0.04 | 3 | 0.01 | 0.28 | 0.84 |
| ABD | 0.07 | 6 | 0.01 | 0.24 | 0.96 |
| ACD | 0.66 | 6 | 0.11 | 2.30 | 0.04 |
| BCD | 0.07 | 2 | 0.03 | 0.69 | 0.50 |
| ABCD | 0.10 | 6 | 0.02 | 0.35 | 0.91 |
| Error | 31.66 | 143 | | | |

| | High light | | | | |
|---|---|---|---|---|---|
| Source of variation | SS | *df* | MS | F | *p* |
| A:Experimental environment | 63.44 | 3 | 21.15 | 758.40 | 0.00 |
| B:Position | 0.07 | 1 | 0.07 | 2.47 | 0.12 |
| C:Previous $CO_2$ history | 1.88 | 1 | 1.88 | 67.39 | 0.00 |
| D:Genotype | 0.44 | 2 | 0.22 | 7.94 | 0.00 |
| Interactions | | | | | |
| AB | 0.02 | 3 | 0.01 | 0.28 | 0.84 |
| AC | 0.85 | 3 | 0.28 | 10.19 | 0.00 |
| AD | 0.85 | 6 | 0.14 | 5.10 | 0.00 |
| BC | 0.10 | 1 | 0.10 | 3.76 | 0.05 |
| BD | 0.10 | 2 | 0.05 | 1.76 | 0.18 |
| CD | 0.25 | 2 | 0.12 | 4.43 | 0.01 |
| ABC | 0.14 | 3 | 0.05 | 1.66 | 0.18 |
| ABD | 0.67 | 6 | 0.11 | 4.01 | 0.00 |
| ACD | 0.63 | 6 | 0.10 | 3.76 | 0.00 |
| BCD | 0.03 | 2 | 0.01 | 0.53 | 0.59 |
| ABCD | 0.34 | 6 | 0.06 | 2.02 | 0.07 |
| Error | 74.00 | 189 | | | |

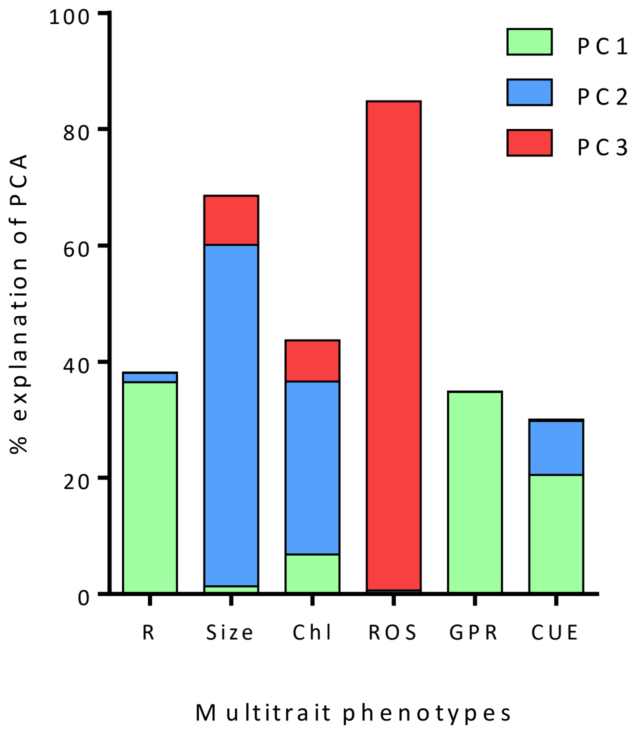


**Figure S1.** Percentage the explanation of each phenotypic trait on the three principal components (PCs) derived from the PCA analysis. The three PCs explained the 77% of the overall variance. Specifically, PC 1 explained 40.71%, PC 2 explained 18.91%, and PC3  explained 17.54%.

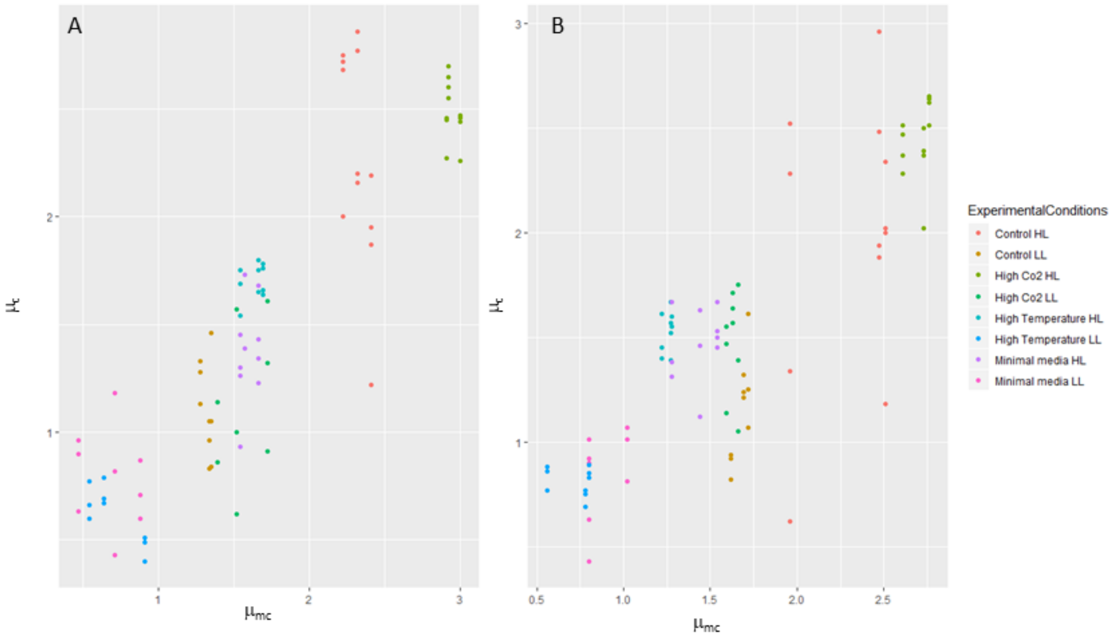


**Figure S2.** Growth in the presence of a non-self population in indirect competition ($\mu_c$) relative to growth in the presence of a self population in indirect competition ($\mu_{mc}$) for ambient-evolved (A) and high-evolved (B) populations.
